# A new paradigm of multiheme cytochrome evolution by grafting and pruning protein modules

**DOI:** 10.1101/2022.02.24.481543

**Authors:** Ricardo Soares, Nazua L. Costa, Catarina M. Paquete, Claudia Andreini, Ricardo O. Louro

## Abstract

Multiheme cytochromes play key roles in diverse biogeochemical cycles, but understanding the origin and evolution of these proteins is a challenge due to their ancient origin and complex structure. Up until now, the evolution of multiheme cytochromes composed by multiple redox modules in a single polypeptide chain was proposed to occur by gene fusion events. In this context, the pentaheme nitrite reductase NrfA and the tetraheme cytochrome *c*_554_ were previously proposed to be at the origin of the extant octa- and nonaheme cytochromes *c* involved in metabolic pathways that contribute to the nitrogen, sulfur and iron biogeochemical cycles by a gene fusion event. Here, we combine structural and character-based phylogenetic analysis with an unbiased root placement method to refine the evolutionary relationships between these multiheme cytochromes. The evidence show that NrfA and cytochrome *c*_554_ belong to different clades, which suggests that these two multiheme cytochromes are products of truncation of ancestral octaheme cytochromes related to extant ONR and MccA, respectively. From our phylogenetic analysis, the last common ancestor is predicted to be an octaheme cytochrome with nitrite reduction ability. Evolution from this octaheme framework led to the great diversity of extant multiheme cytochromes analised here by pruning and grafting of protein modules and hemes. By shedding light into the evolution of multiheme cytochromes that intervene in different biogeochemical cycles, this work contributes to our understanding about the interplay between biology and geochemistry across large time scales in the history of Earth.

## Introduction

Multiheme cytochromes *c* (MHC) catalyze diverse chemical reactions in prokaryotes providing them with a remarkable metabolic versatility (Simon et al. 2011; Bewley et al. 2013; Deng et al. 2018). These metalloproteins exploit the redox, spin and acid-base properties of the heme cofactors to perform chemical reactions that are pivotal to many biogeochemical cycles, including those of nitrogen, sulfur and iron (Pereira and Xavier 2006; Mayfield et al. 2011; Paquete et al. 2019). Some MHC families share sequence and structural similarities that unequivocally reflect a common ancestral origin (Sharma et al. 2010). Understanding their evolution has been a challenge due to the low preserved amino acid sequence and limited available 3D structures that are better preserved than sequence across large timescales (Chothia and Lesk 1986; Illergård et al. 2009; Ingles-Prieto et al. 2013). Nevertheless, the current paradigm for the emergence of the different known MHC follows the proposed fusion of redox modules, going from simple MHC to more complex and containing more heme cofactors per polypeptide chain. This comes from the observation of the presence of repetitive redox modules in MHC that has enabled to relate different MHC families (Roldán et al. 1998; Arménia Carrondo et al. 2006; Santos-Silva et al. 2007; Clarke et al. 2011; Pokkuluri et al. 2011; Pereira et al. 2017; Edwards et al. 2020).

In the case of the pentaheme nitrite reductase (NrfA) and the octaheme hydroxylamine oxidoreductase (HAO), structural similarities of the heme-core and of the three interface-forming helices has long been pointed out as a sign of the common origin between these two MHC that have distinct functions in the nitrogen cycle (Igarashi et al. 1997; Einsle et al. 1999). Octaheme nitrite reductases (ONR) later discovered, showed even more similarities with NrfA with respect to its structure and function (Polyakov et al. 2009; Tikhonova et al. 2012). Cytochrome *c*_554 (_cyt *c*_554)_ that is unrelated to NrfA, also showed similarities with HAO regarding the heme-core structure (Iverson et al. 1998), suggesting that diverse evolutionary mechanisms took place for the emergence of these proteins. In all of these proteins, catalysis takes place at a heme with one open coordination in the iron for access of the substrate.

Using phylogenetics and placing NrfA at the root, an ancestral ONR was proposed to be an intermediary for the evolution of NrfA to the presumably more recent HAO family of proteins (Klotz et al. 2008). In that proposal, an unknown triheme cytochrome fused with NrfA to originate ONR. That simple evolutionary scheme required revision as more homologous MHC were characterized. For example, the structure of the copper sulfite reductase MccA (Hermann et al. 2015), which also reduces nitrite, and the octaheme nitrite reductase IhOCC (Parey et al. 2016) revealed similarities with NrfA, ONR and HAO. However, unlike ONR and HAO, the catalytic heme of MccA is located in the N-terminal region (Hermann et al. 2015). This feature is also found in the octaheme tetrathionate reductase (OTR) (Mowat et al. 2004) that was proposed to have diverged from NrfA via a different route based on the lack of the interface-forming helices characteristic of HAO and ONR (Klotz et al. 2008). More recently, the structure of the cell-surface nonaheme OcwA from the electroactive bacterium *Thermincola potens* JR was determined (Costa et al. 2019). This protein, which has iron-oxide reductase activity *in vitro* in agreement with its proposed physiological role, did not reveal structural similarities with other structurally characterized cell-surface iron reductases (Edwards et al. 2012; Edwards et al. 2015; Wang et al. 2019; Edwards et al. 2020). By contrast, OcwA showed similarities with NrfA at the C-terminus and cyt *c*_554_ at the N-terminus, conserving the catalytic hemes of both proteins. These observations led to the proposal that an ancestral OcwA-like protein could represent the evolutionary link between NrfA and cytochrome *c*_554_ with the different octaheme cytochromes (Costa et al. 2019). In this scenario, OcwA would have originated from a gene fusion event between ancestral NrfA and cyt *c*_554_. Subsequent loss of one of the two catalytic hemes and diversification would have led to the different extant octaheme cytochromes (Costa et al. 2019). However, these previous studies (Klotz et al. 2008; Costa et al. 2019) lacked a phylogenetic analysis with a combination of structural and character (sequence) information, and an unbiased root placement method. Here, we used these criteria combined with minimal functional site characterization of the hemes to refine the previous evolutionary proposals. Our phylogenetic analysis revealed that cyt *c*_554_ and NrfA are products of a truncation event from different octaheme cytochromes and that the last common ancestor (LCA) is inferred to be an octaheme cytochrome able to reduce nitrite.

## Results

### Identification of the homologous group of MHC

The MHC analyzed in this work (table 1) are involved in diverse metabolic pathways. NrfA, ONR, IhOCC, MccA, OTR are involved in dissimilatory nitrite and/or sulfite and/or tetrathionate reduction. Cyt *c*_554_, HAO and hydrazine dehydrogenase (HDH) are involved in ammonia oxidation reactions. HAO oxidizes hydroxylamine aerobically; HDH oxidizes hydrazine anaerobically; cyt *c*_554_ functions as an electron transfer protein from HAO to membrane-bound cytochrome *c*_552_ (for a review see Paquete et al. (2019)). OcwA is involved in dissimilatory iron reduction by extracellular electron transfer (Carlson et al. 2012; Costa et al. 2019). Nevertheless, all of them display nitrite or nitric oxide reductase activity *in-vitro* (Costa et al. 2019; Paquete et al. 2019). Sequence and structural similarities allowed the assignment of the recently identified undecaheme OmhA (Gavrilov et al. 2021) as belonging to this homologous group (table 1). This protein was isolated from the S-layer of the thermophilic and Gram-positive bacterium *Carboxydothermus ferrireducens* (Gavrilov et al. 2012). OmhA shares 29.5% semi-global sequence identity and its structure has a C-α root mean square deviation (RMSD) of 2.45Å (PyMOL, Schrödinger, Inc.) with OcwA. The amino acid sequence of OmhA contains an extra N-terminal extension harboring two extra hemes in comparison with OcwA (supplementary fig. S1). Sequence and structural searches did not show any homologous MHC to this small region.

**Table 1.**
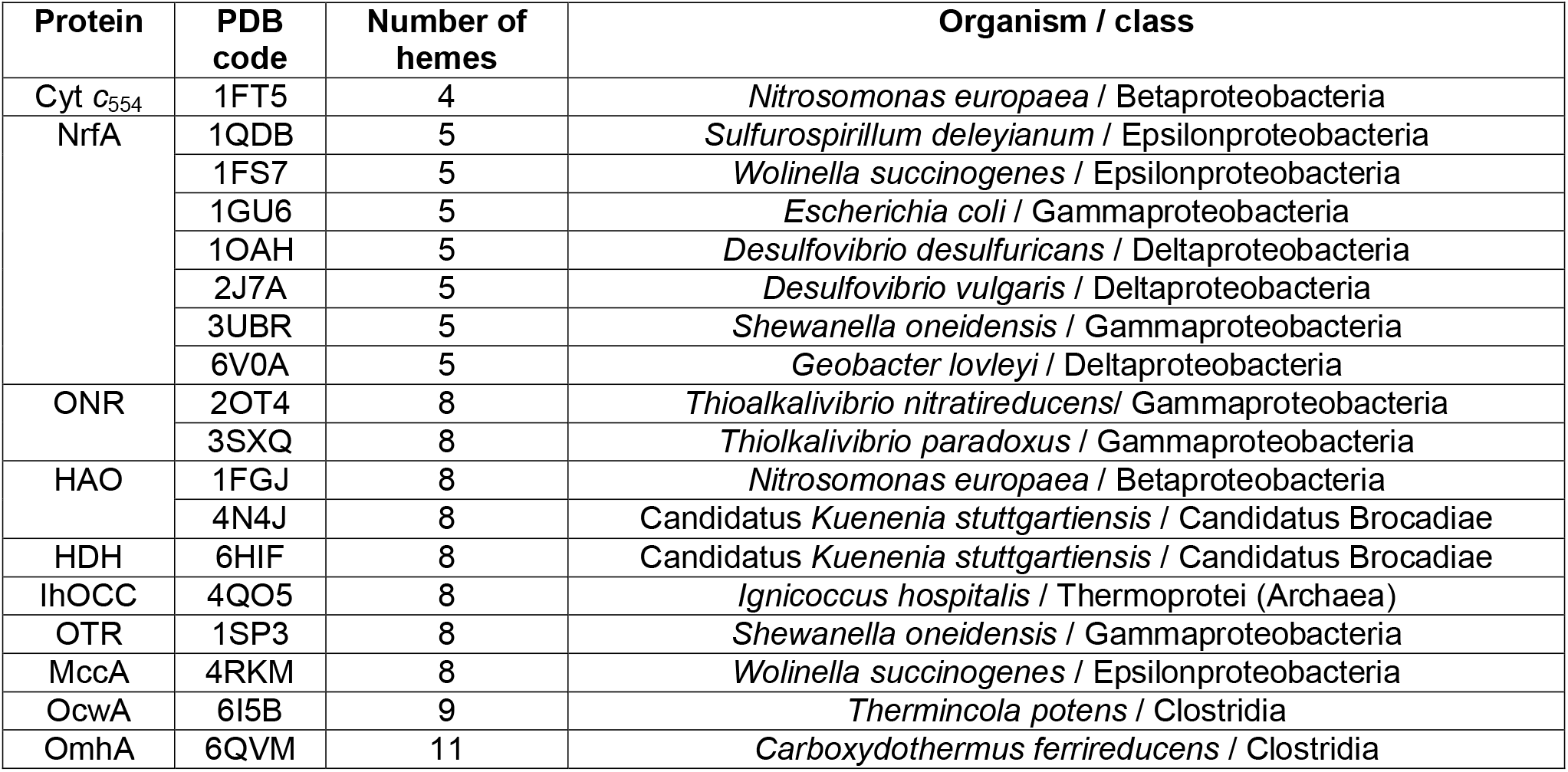
List of the homologous MHC. For each MHC, the PDB code of the 3D structure, the number of hemes and the source organism and the taxonomic class are indicated.

The MHC listed in table 1 share one or several structurally homologous features (fig. 1). The exception is cyt *c*_554_ and NrfA that lack similarity between each other as these two cytochromes align at opposing ends (N- and C-terminal, respectively) of the octa-, nona- and undecaheme cytochromes. Based on this observation, cyt *c*_554_ and NrfA are used to define N- and C-terminal modules of the MHC analysed in this work (fig. 1), respectively. All other MHC share a similar heme-core arrangement with three or more hemes within these modules displaying good structural superimposition. One key difference is the presence of one or two catalytic hemes. NrfA, ONR, HAO, HDH and IhOCC contain the catalytic heme within the “C-terminal module”, while cyt *c*_554_, MccA and OTR contain the catalytic heme within the “N-terminal module” (fig. 1). OcwA and OmhA are currently the only proteins that preserve both catalytic hemes. Another notable observation is the presence or absence of the three C-terminal oligomerization-forming helices that are absent in cyt *c*_554_ (lacks the C-terminal module) and in OTR (fig. 1). The distal axial ligands’ position is conserved in all MHC except in OTR where the distal ligands of hemes 1 and 7 are found in different relative positions in the amino acid sequence (supplementary fig. S2).

**Fig. 1.**
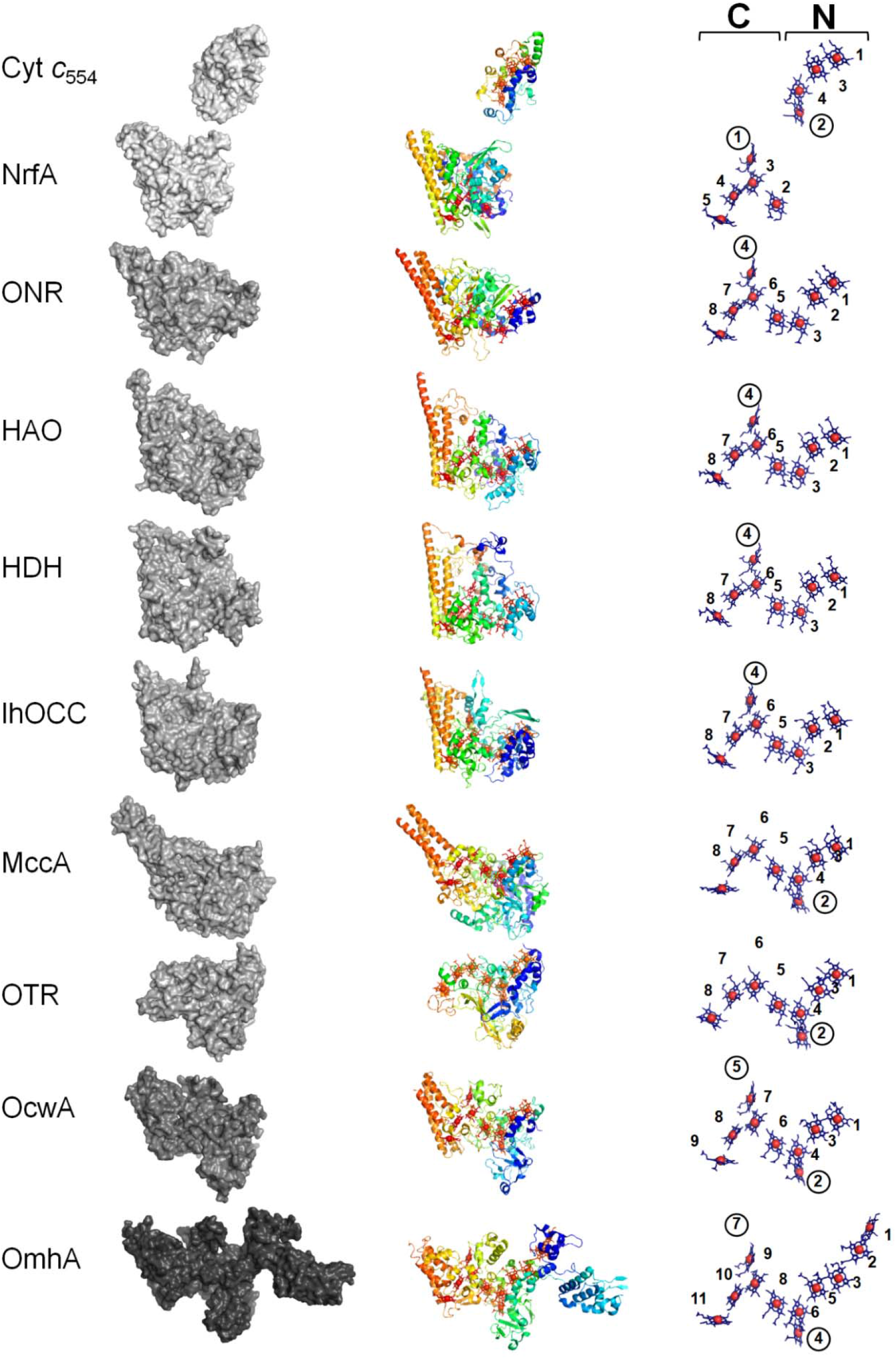
Aligned structures for the representative MHC of the three main clusters. PDB structures used were Cyt *c*_554_, 1FT5_A; NrfA, 1QDB_A; ONR, 2OT4_A; HAO, 1FGJ_A; HDH, 6HIF_A; IhOCC, 4QO5_A; OTR, 1SP3_A; MccA, 4RKM_A; OcwA, 6I5B_A; OmhA, 6QVM_A. Surface, cartoon and heme core representations are depicted. Cartoon representations are rainbow colored from N-(right) to C-(left) terminal. Heme-core representation contains heme position number and catalytic hemes (pentacoordinated) are identified by a circle. The C- and N-terminal modules are indicated by the respective letters.

### Comparison of the backbone trace and heme-core architecture

In order to assess the structural distance within these MHC, the backbone and heme-core arrangements were compared. As shown by the network representation in fig. 2 (corresponding distance matrices are available in supplementary table 1 – 4) OTR is significantly different from all other MHC. This protein appears as an outlier with respect both to the backbone and heme-core structural comparisons (fig. 2). Based on the backbone trace and heme-core architecture, the remaining MHC clustered in three distinct clades: HAO, HDH and IhOCC (clade1); NrfA and ONR (clade 2); cyt *c*_554_, MccA, OcwA and OmhA (clade 3). In clades 1 and 2, intra-cluster structural distances are globally shorter than in clade 3, which did not always meet the selected threshold for node (structure) connection (grey lines in fig. 2). Nevertheless, clade 3 MHC were consistently found together when compared with the rest of the MHC (fig. 2), with the exception of the backbone trace comparisons at the C-terminal where clade 2 and 3 are close together by the short distance between OcwA and one NrfA structure (PDB entry 3UBR) (fig. 2B). Within clade 3, the cell-surface MHC OcwA and OmhA presented the shortest distances, with the exception for the structural comparisons of the heme-core arrangement at the C-terminal (fig. 2D; supplementary table 4). In this case, OcwA and MccA pair showed the shortest distances.

**Fig. 2.**
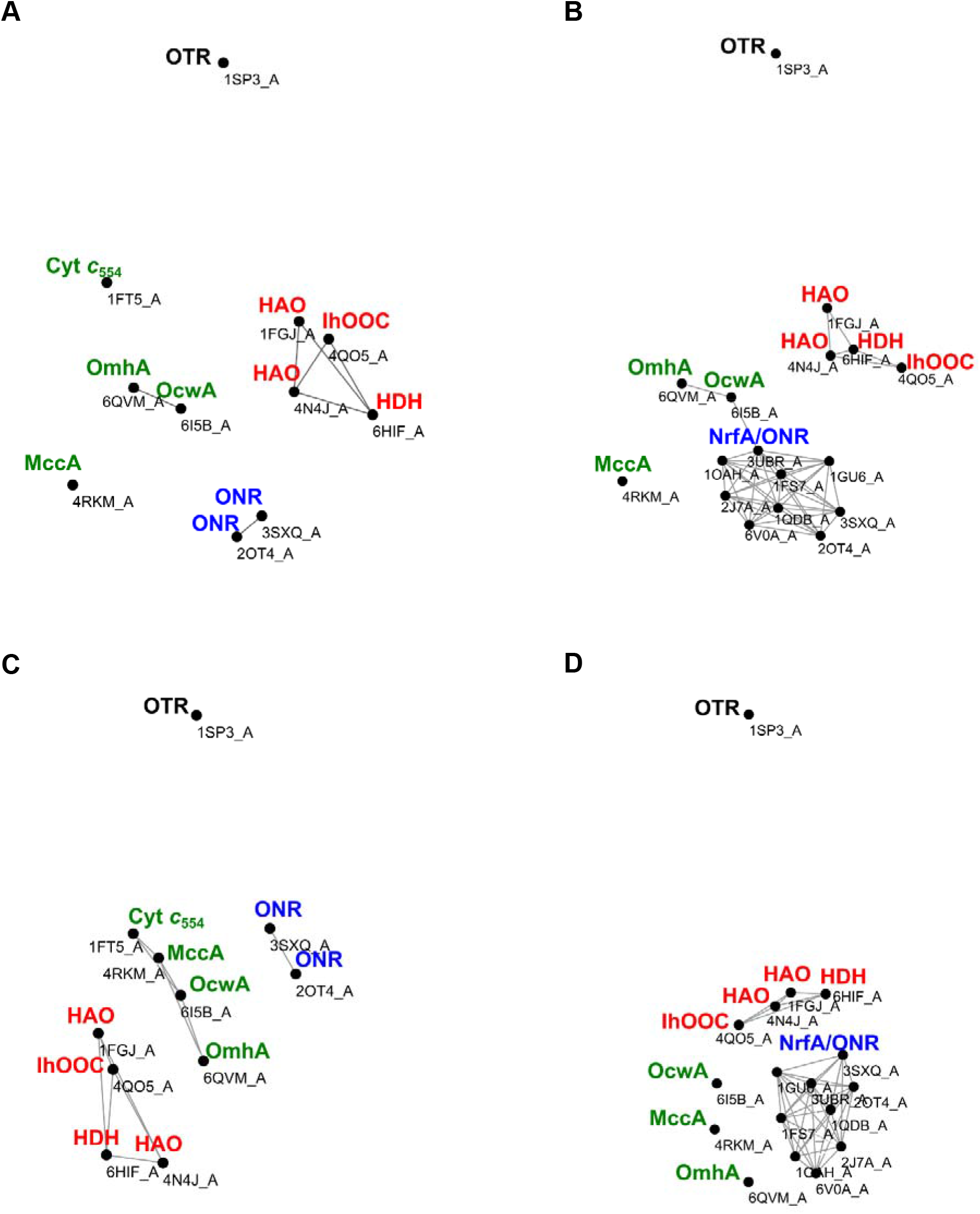
Distance network analysis using structural information. Structural comparisons were performed using distance comparisons of the backbone structure at the N-terminal (*A*) and C-terminal (*B*) and of the heme core at the N-terminal (*C*) and C-terminal (*D*). Each node (black dot) represents one structure. Grey lines connect structures with distances equal or less than 3.5 and 0.75 ångströms for the backbone and heme core structures, respectively.

### Comparison of the heme centers as minimal functional sites

Minimal functional sites in metalloproteins (MFSs) are portions of the 3D structure that focus on the region around the metal site(s). MFSs of heme centers were extracted from representative structures of the MHC listed in table 1, compared all versus all with the MetalS^2^ tool (Andreini et al. 2013), and clustered into groups of structurally similar MFSs. This analysis resulted in 10 groups, comprising 72 MFSs out of the 77 included in the analysis, which were arbitrarily named group A (26 MFSs), group B (11 MFSs), group C (9 MFSs), group D (6 MFSs), group E (5 MFSs), group F (4 MFSs), group G (4 MFSs), group H (3 MFSs), group I (2 MFSs), and group J (2 MFSs). Table 2 shows the groups to which each heme center in the MHC belongs. Since MFSs describe the local environment around a metal cofactor, each group of MFSs identifies a set of heme centers in MHC with common structural features, which for the sake of simplicity we refer to as a heme “type”. Observation of table 2 confirms that (i) OTR represents an outlier with respect to all the other MHC, since only the heme centers at positions 1 and 8 are of a type shared by other MHC; and (ii) HAO, HDH and IhOCC represent a clear subgroup (clade 1 above), since all of them have the same pattern of heme types, which includes three centers (at positions 1, 2, and 5) different from all the other MHC. The relationships between the other MHC (ONR, NrfA, OcwA, OmhA, MccA, and cyt c_554_) are not clearly defined with this analysis, since the differences are mainly related with absence of specific hemes or the presence of heme types that are not clustered.

**Table 2.** MFS groups by heme position in the amino acid sequence within each MHC. For each MHC, the PDB code of the 3D structure, the position of hemes in the sequence and the group of MFSs to which each heme center belongs are indicated. “Other” means that the heme center was not clustered in any group.

|  | **-1** | **0** | **1** | **1bis** | **2** | **3** | **4** | **5** | **6** | **7** | **8** |
| --- | --- | --- | --- | --- | --- | --- | --- | --- | --- | --- | --- |
| HAO |  |  | H |  | B | A | C | G | A | B | A |
| HDH |  |  | H |  | B | A | C | G | A | B | A |
| IhOCC |  |  | H |  | B | A | C | G | A | B | A |
| ONR |  |  | E |  | Other | A | C | D | A | B | A |
| NrfA |  |  |  |  |  |  | C | D | A | B | A |
| OcwA |  |  | E | Other | F | A | C | D | A | B | A |
| OmhA | D | Other | E | C | F | A | C | D | A | B | A |
| MccA |  |  | A | Other | F | A |  | D | A | B | A |
| Cyt *C*<sub>554</sub> |  |  | E | C | F | A |  |  |  |  |  |
| OTR |  |  | E | Other | G | I |  | J | I | J | A |

Overall, the results of the analysis of MFSs is consistent with those obtained for the **Comparison of the backbone trace and heme-core architecture**, which found HAO and IhOCC clustered in clade 1 and the remaining MHC divided in clades 2 and 3 (fig. 2).

### Character-based phylogeny

A total of 5700 sequences were collected from NCBI RefSeq database (supplementary table 5). These correspond to highly homologous sequences to at least one of the reference MHC listed in table 1. These sequences ranged from 23.1 to 100 % of global identity. Globally the diversity of prokaryotes harboring homologous MHC was greatly expanded from those whose MHC have been structurally characterized (table 1). A total of 22 phyla and two domains (Bacteria and Archaea) were found in this analysis (supplementary table 5). NrfA was the most represented family within the database in terms of number of sequences and diversity, while the cell-surface MHC (OcwA and OmhA) were very poorly represented. Since OTR was shown to be an outlier in the structural comparisons (see **Backbone trace, heme-core architecture and minimal functional site comparisons**) it was not included in this analysis. HAO and HDH homologues greatly overlapped as HAO and HDH from *Kuenenia stuttgartiensis* are phylogenetically closer than HAO from *Nitrosomonas europaea*. In this sense, this group was joined hereon (HAO/HDH).

Phylogenetic inference was performed using multiple sequence alignments (MSA) either containing NrfA (NrfA+) or cyt *c*_554_ (cyt *c*_554_+) along with the rest of the MHC, as these cytochromes align at opposing ends of the sequence of the remaining MHC. Reconstructed phylogenies consistently indicated the presence of three main monophyletic clades composed by HAO/HDH and IhOCC (clade 1), NrfA and ONR (clade 2) and OcwA, OmhA, MccA and Cyt *c*_554_ (clade 3) (fig. 3A–D). All monophyletic proposed clades are supported with SH-aLTR/Bootstrap or posterior probabilities above 70%. For clades 1 and 2 these statistical support values are above 90%. Within the last clade the cell-surface MHC were clustered together (OcwA and OmhA). Likewise, MccA and cyt *c*_554_ were clustered together within this clade. Minimal ancestor deviation rooting analysis (Tria et al. 2017) consistently placed the root between clade 1 and the junction of clades 2 and 3.

**Fig. 3.**
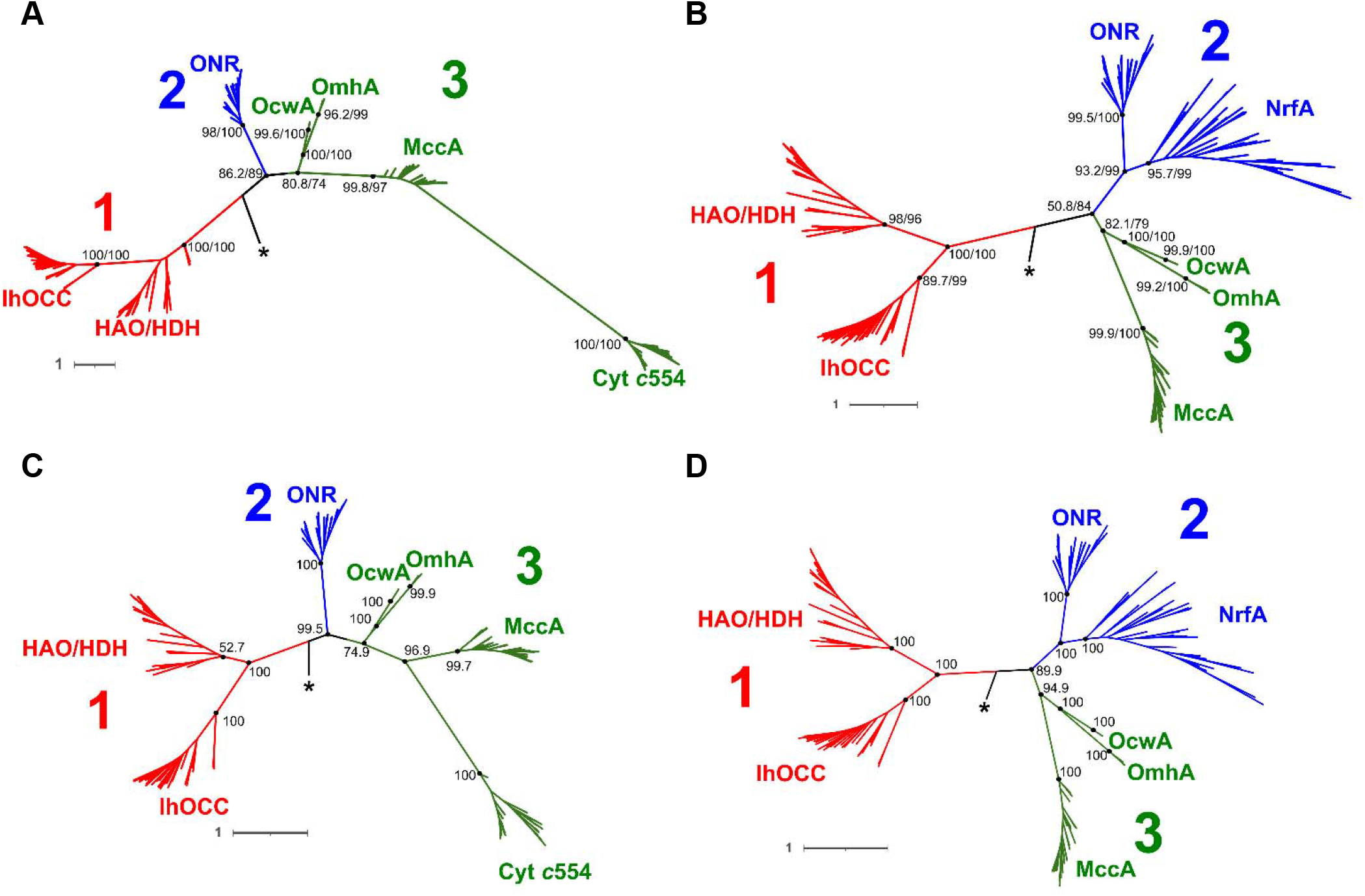
Character-based phylogenetic analysis for the homologous MHC. Maximum likelihood phylogenetic trees based on cyt *c*_554_+ (*A*) and NrfA+ (*B*) multiple sequence alignments. Bayesian phylogenetic trees based on cyt *c*_554_+ (*C*) and NrfA+ (*D*) multiple sequence alignments. Protein sequences fell in three main clades according with their tree positioning and are represented in different colors (red, blue and green). Asterisks indicates the root placement according with the minimal ancestor deviation method (Tria et al. 2017). Bootstrap/SH-aLTR (ML trees) or posterior probabilities (Bayesian trees) confidence percentages values are presented near each node for the major splits. Tree scale represents number of substitutions per site.

Mismatch between the phylogenetic position of some sequences and the taxonomic classification of the organisms from which these sequences derive were observed across all clades analyzed (supplementary fig. S3, 4 and 5). This suggests widespread occurrence of horizontal gene transfer (HGT) for these MHC. Notably, OcwA and OmhA sequences only presented vertical transfer in the current dataset. However, this may be a consequence of the poor representation of these two MHC in the public database analyzed. It contains only 2 and 4 highly homologous sequences, respectively, and from bacteria belonging the same genus, *Thermincola* and *Carboxydothermus*, respectively (supplementary table 5).

### Conservation of critical residues for function

In order to gain further insights about the emergence of the unique features that differentiate these MHC, the conservation of critical residues for function were searched and compared within each MHC clade (fig. 4). The ability to oxidize hydroxylamine or hydrazine appears to have a relatively recent origin within clade 1, judging by the branching points of the homologous sequences that conserve the tyrosine residue at the crosslink position (supplementary fig. S6). In clade 2 all ONRs and NrfAs contain the typical known features for nitrite reduction, namely the lysine as the proximal ligand of the catalytic heme and the catalytic tyrosine, histidine and arginine, which correspond to K131, Y217, H282 and R113 in *Sulfurospirillum deleyianum* (PDB entry 1QDB_A)), respectively. The only exception are the NrfA variants without the lysine proximal ligand of the catalytic heme (supplementary fig. S7) that appear to have emerged recently within clade 2. Clade 3 is more diversified in terms of function and structure than clade 1 and 2. Nevertheless, the identified residues that are important for catalysis are all conserved in OcwA and MccA, which have catalytic residues assigned in their structures. All OcwA and OmhA sequences conserve the catalytic histidines that are localized near the catalytic hemes. These correspond to H282 and H381 in *Thermincola potens* JR (PDB entry 6I5B_A). OcwA and OmhA conserve also two extra histidines nearby the first and second catalytic hemes (positions 1bis and 4 in table 2), H281 and H380 in *Thermincola potens* JR (PDB entry 6I5B_A). *Thermincola potens* JR H380 is conserved in all OcwA and OmhA sequences, while *Thermincola potens* JR H281 is conserved in all OcwA sequences and in half of the OmhA sequences. All MccA sequences conserve the known residues that are important for catalysis, namely Y123, K208, Y285, Y301, R366, K393, C399 and C495 in *Wolinella succinogenes* (PDB entry 4RKM_A). The eighth heme (position 8 in table 2) has the uncommon CX_15_CH motif (Hermann et al. 2015). In addition, we found also the presence of the CX_17_CH heme-binding motif for the eighth heme of the MccA sequences in our dataset.

**Fig. 4.**
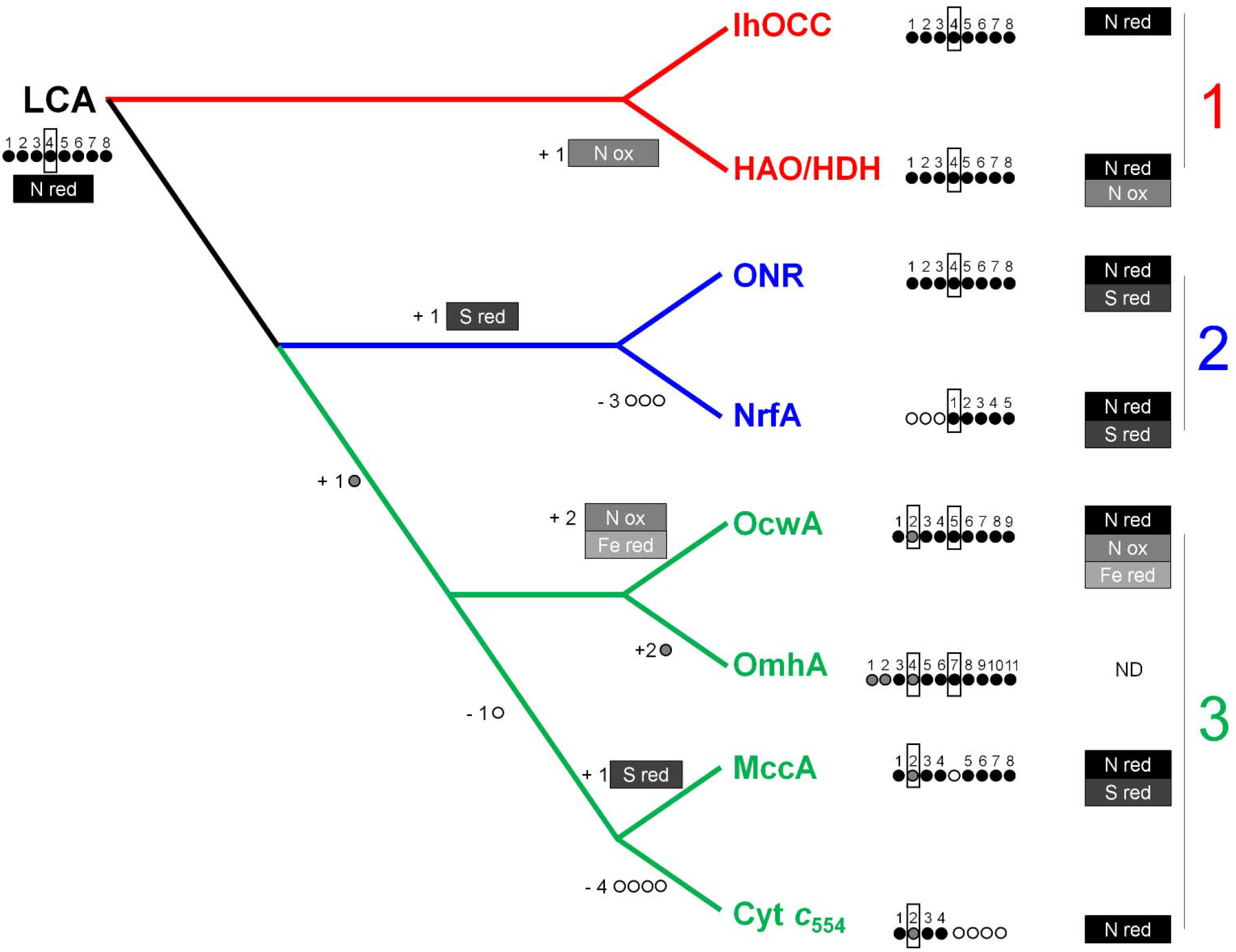
Cladogram with the most parsimonious solution for the evolution of the MHC after phylogenetic analysis. Circles and rectangles represent the hemes and catalytic activities, respectively, for each MHC. Dark circles and rectangles represent hemes and activities inherited from the last common ancestor (LCA) while grey and white represents heme or activity gain and loss, respectively. Gains and losses of hemes and activities from the LCA to extant MHC are depicted near each respective position in the cladogram. Catalytic hemes are marked within black rectangles. ND represents no data.

## Discussion

The dissection of the origin and evolution of MHC has attracted significant scientific interest but has also been characterized by numerous challenges due to the small sequence similarity that was preserved among these proteins along time. Nevertheless, a great number of MHC have been identified as homologues based on structural similarities. The timeline for their emergence is typically thought to progress by fusion of redox modules towards more complex MHC. These can reach up to 16 hemes for structurally characterized MHC, but genes coding putative MHC containing as much as 113 hemes per polypeptide chain have been reported (Roldán et al. 1998; Arménia Carrondo et al. 2006; Santos-Silva et al. 2007; Clarke et al. 2011; Pokkuluri et al. 2011; Pereira et al. 2017; Edwards et al. 2020; Leu et al. 2020). Towards understanding the evolution of MHC we focused our attention on a group of MHC that is involved in the biogeochemical cycles of iron, nitrogen and sulfur.

In addition to gene fusion, gene fission is also a well-established mechanism for diversity of domain combination in all domains of life (Kummerfeld and Teichmann 2005; Pasek et al. 2006; Weiner and Bornberg-Bauer 2006). However, only gene fusion was considered while proposing the origin of the octa and nonaheme cytochromes by NrfA and cyt *c*_554_ (Klotz et al. 2008; Costa et al. 2019). Indeed the previous proposals for MHC evolution were biased towards a fusion event (Klotz et al. 2008; Costa et al. 2019). By performing a thorough phylogenetic inference with both structural and sequence data we now show that NrfA and cyt *c*_554_ were likely formed by truncation of octaheme cytochromes from different clades. Phylogenetic analysis revealed three main clades composed by HAO/HDH and IhOCC (clade 1), NrfA and ONR (clade 2) and OcwA, OmhA, MccA and cyt *c*_554_ (clade 3). NrfA and cyt *c*_554_ belong to different clades and consequently, the MHC that are phylogenetically closer to NrfA are not the same for cyt *c*_554_. The closest homologue of NrfA is ONR, while for cyt *c*_554_ the closest homologue is MccA. This implies that a fusion event between NrfA and cyt *c*_554_ was not responsible for OcwA origin.

Although previous studies presented unrooted trees (Bergmann et al. 2005; Klotz et al. 2008), NrfA was chosen as the implicit root for those phylogenetic analyses, based on the premise that nitrate/nitrite ammonification is the most ancestral reaction when compared with ammonia oxidation reactions. However, for a global cycle dating perspective, isotope signatures cannot at present, discriminate between all of the different reactions within the nitrogen cycle (Stüeken et al. 2016). Moreover, since those studies other MHC capable of nitrite reduction besides NrfA were identified (e.g. IhOCC (Parey et al. 2016)) and dissimilatory iron reduction is also considered to be a very early form of bioenergetic metabolism (Vargas et al. 1998; Johnson et al. 2008). Attempts at a rigorous assessment of relative dating using taxonomics in order to solve the root placement is complicated by the evidence that most of these MHC have undergone considerable HGT, which was detected in the present analysis. Likewise, HGT of MHC has been extensively reported in the literature, including for NrfA (Welsh et al. 2014), HAO (Bergmann et al. 2005) and several MHC involved in extracellular electron transfer, such as MtrCAB and OmcA from *Shewanella* spp. (Zhong et al. 2018; Baker et al. 2022).

Outgroup rooting is by far the most used method to infer the root position in a given phylogenetic tree but it is unsuitable when the ingroup is already very divergent and no appropriate outgroup is available. Considering this, we used Minimal Ancestral Deviation method for rooting our trees (Tria et al. 2017). This method is an updated version of the midpoint rooting method by enabling perturbations to the molecular clock and therefore heterotachy. This analysis placed the root between clade 1 and the junction of clades 2 and 3. Using this information we can conclude that clade 1 is the most ancient, as it represents the basal clade from the root position, followed by clade 2 and clade 3.

Using the most parsimonious solution for inferring ancestral states, the LCA is inferred to be an octaheme cytochrome that was able to reduce nitrite. In the timeline that we propose (fig. 4), clade 1 was the first to branch into HAO/HDH and IhOCC, both having the same heme-core architecture of the LCA. The catalytic heme of HAO/HDH is a P460 heme and most of the sequences collected for HAO/HDH conserve the tyrosine residue at crosslinking position of the representative structures (supplementary fig. 6). Contrarily, the P460 heme is absent in IhOCC and it is predicted to be absent also in the LCA. In this scenario, the P460 heme evolved *de novo* in the HAO/HDH subclade. Indeed, an HAO family protein was recently isolated from *K. stuttgartiensis* and showed no hydroxylamine oxidation activity, while presenting nitrite reduction to nitric oxide (Ferousi et al. 2021). The second branching clade was clade 2, that latter diversified into ONR that conserved the overall heme-core arrangement of the LCA, and NrfA that further lost the three N-terminal hemes. The lysine residue as the proximal ligand of the catalytic heme seems to be an innovation of this clade, conserved in all ONR sequences. By contrast, not all sequences that were collected for NrfA conserve this lysine residue even though no structure is available for these variants. Clade 3 appeared more recently by the addition of a second catalytic heme (position 1bis in table 2). OcwA and OmhA still conserve also the first catalytic heme (position 4 in table 2) that is common to the two more ancient clades (1 and 2), and OmhA gained two extra hemes at the N-terminal. By contrast, MccA and cyt *c*_554_ have lost the first catalytic heme (position 4 in table 2), and cyt *c*_554_ further lost the last four C-terminal hemes. Inference that the LCA was an octaheme cytochrome and that loss of hemes took place, was previously proposed by Kern et al. (2011). However, figure 1 shows that the sequence alignments used by Kern et al. (2011), are not congruent with the respective 3D structure alignments. Kern et al. (2011) aligned the catalytic hemes of MccA and cyt *c*_554_ (position 1bis in table 2) with the ones of NrfA, ONR and HAO (position 4 in table 2), which results in an 180 degree y axis misalignment from the aligned protein structures of these two groups presented in figure 1. Two consequences arise from this: the tree topology artificially exacerbates the distance between these two groups and by proposing an alignment of NrfA with cyt *c*_554_, Kern et al. (2011) failed to realize the existence of redox modules, that is clearly apparent in figure 1.

The cladogram of fig. 4 provides additional support to the analysis performed here given that it recapitulates the patterns of heme “types”, as described by MFSs, observed in extant MHC (supplementary fig. S8). In this scenario, clade 1 conserved the local heme environments from the LCA, while clade 2 and 3 underwent successive modifications. The common branching of clade 2 and 3 from clade 1 involved the modification of the two first hemes (position 1 and 2 in table 2). The subsequent origin of clade 3 involved the addition of a new catalytic heme (position 1bis in table 2) that had similar characteristics of the first catalytic heme (position 4 in table 2) and further modification of the heme at position 2 (table 2). Diversification of clade 3 involved further modifications of the second catalytic heme (OcwA and MccA) to new specific heme types (position 1bis in table 2). Addition of two extra N-terminal hemes (positions −1 and 0 in table 2) for OmhA and modification of the heme at position 1 to the same type of positions 3, 6 and 8 (table 2) in the case of MccA also led to the diversification of the heme patterns of clade 3.

## Conclusion

MHC were abundantly employed by nature in the development of multiple biogeochemical cycles across large time scales, which were of high importance for the colonization of all extant ecological niches of life on earth. Previous proposals for the evolution of MHC have been biased towards gene fusion events (Roldán et al. 1998; Arménia Carrondo et al. 2006; Santos-Silva et al. 2007; Clarke et al. 2011; Pokkuluri et al. 2011; Pereira et al. 2017; Edwards et al. 2020). Our study changes this perspective by showing that fission of heterogeneous redox modules also drives the evolution of MHC and should be *a priori* equally considered. It shows that NrfA and cyt *c*_554_ likely resulted from truncation events of an ancestral ONR and MccA, respectively, and that the common ancestor of the MHC analysed here was likely an octaheme cytochrome similar to extant IhOCC with nitrite reductase activity. Evolution from this ancestral octaheme cytochrome included pruning and grafting of heme-binding polypeptide modules, which led to the emergence of extant MHC that catalyze very distinct reactions within the nitrogen, sulfur and iron biogeochemical cycles. Altogether, this work opens a new perspective in our understanding about the evolution of MHC and their changing role in the interplay between biology and geochemistry across large timescales.

## Materials and Methods

### Backbone and heme-core comparisons

Protein structures were retrieved from the PDB database (table 1). Chain A of each protein was selected for subsequent structural comparisons. The data was divided in two subsets for analysis, one including all MHC except NrfA and another including all MHC except cyt *c*_554_, as these align at opposing ends of the octa-, nona- and undecaheme MHC. Heme-core comparisons were performed using PyMOL Molecular Graphics System (version 2.0 Schrödinger, LLC). The structural positions of the 33 non-mobile atoms of the porphyrin ring were selected for each heme. For the analysis including cyt *c*_554_ the three common N-terminal hemes to all MHC except NrfA were selected (positions 1, 2 and 3 in table 2). For the analysis including NrfA, the four common C-terminal hemes to all MHC except cyt *c*_554_ were selected (positions 5, 6, 7 and 8 in table 2). Pair-fit function was used to align the corresponding positions of each pair. Pairwise RMSD values were used to construct a distance matrix that was then fed into Cytoscape 3.7.1 to construct networks based on structural distance (Shannon et al. 2003). Representation was generated using the Prefuse force directed layout algorithm with 1 - normalized weight values option and 1000 iterations. Backbone structure of the MHC was compared using mTM-align (Dong, Pan, et al. 2018; Dong, Peng, et al. 2018). From the multiple structure alignment generated, pairwise RMSD were retrieved and used to build a matrix that was then used as input data for network representation in Cytoscape, in a similar procedure to that performed for the heme-core structural comparisons.

### Minimal functional sites analysis

MFSs were extracted for each heme in the PDB structures listed in fig. 1 and compared using the MetalS^2^ tool (Andreini et al. 2013). The MFSs were then clustered by average linkage clustering using a threshold value of 2.25, which has been previously shown to indicate significant structural similarity between sites (Rosato et al. 2016). This procedure resulted in the clustering of 69 out of a total of 77 MFSs. A further three MFSs were included in clusters as all their similarity scores below 2.50 (Valasatava et al. 2015) were associated with MFSs all belonging to the same cluster.

### Sequence collection and clustering

Sequences were retrieved from NCBI Reference Sequence database (RefSeq) (O’Leary et al. 2016) using Blastp algorithm (Altschul et al. 1990). Sequences from all the proteins whose structure has been determined were used as queries (access date: 2021/15/09). An *e*-value of 1^-50^ was selected as a threshold. Protein sequences were filtered accordingly with the expected number of heme-binding motifs (e.g. 5 heme-binding motifs for NrfAs). The common heme-binding motif (CX_2_CH) and also other less common heme-binding motifs (CX_2_CK, CX_3_CH, CX_4_CH, CX_11_CH, CX_15_CH and CX_17_CH) were used in the filtering steps. CD-HIT (Li et al. 2001; Huang et al. 2010) was used to cluster highly homologous sequences in order to reduce the size of the dataset but maintain the overall diversity. A total of 25 representative sequences without reference sequences (that were previously used as queries) were selected for each group. Reference sequences that contain 3D structures were added separately.

Collected sequences for each protein group were aligned using MAFFT 7 (Katoh and Standley 2013). Using MEGA 7 platform (Kumar et al. 2016), overall mean distances were individually computed using the *p*-distance model (Nei and Kumar 2000), uniform rates and pair-wise deletion methods. Standard errors were calculated using Bootstrap method with 1000 replications (supplementary table 5).

### Sequence alignments

For phylogenetic analyses, two multiple sequence alignments (MSA) were performed. One MSA contained protein sequences for all targeted MHC except NrfA (*c*_554_+ MSA) and the second MSA contained all MHC except cyt *c*_554_ (NrfA+ MSA). Globally, protein sequences that included examples of reported 3D structures were aligned using PROMALS 3D (Pei et al. 2008) with default parameters. This program integrates sequence information derived from predicted secondary structure, profile-profile Hidden Markov Models and structural information derived from sequence-structure and structure-structure alignments as constraints for consistency-based progressive alignments. In addition, user-defined constraints of alignable heme-binding motifs and other important and identified structural elements, e.g. distal axial ligands and active site residues were used as anchors for the alignment. This MSA was used as a structural reference alignment in MAFFT 7 (Katoh and Standley 2013) to align the sequences retrieved from NCBI (see **Sequence collection and clustering**). The L-INS-I algorithm was used with the ‘leave gappy regions’ option. Resulting MSA was inspected in MEGA 7 and manual refinement was performed when necessary. Low-quality positions containing more than 75% gaps were filtered with trimAl version 1.3 (Capella-Gutiérrez et al. 2009) within the Phylemon 2.0 platform (Sánchez et al. 2011). For the pairwise sequence alignment of OcwA and OmhA, the Needleman–Wunsch algorithm (Needleman and Wunsch 1970) was used and represented in ESPript 3.0 (Robert and Gouet 2014).

### Phylogenetic analysis

Since NrfA and cyt *c*_554_ align at opposing ends of the octa-, nona- and undecaheme MHC, phylogenetic inference was performed using two MSA in separate (cyt *c*_554_+ and NrfA+ MSA) (see **Sequence alignments**) generating two phylogenetic trees. Each tree was reconstructed using maximum likelihood and Bayesian methods. Cyt *c*_554_+ MSA contained 749 positions and NrfA+ MSA contained 793 positions. For maximum likelihood inference, phylogeny was reconstructed using IQ-tree (version 2.1.2 COVID-edition built Oct 22 2020) (Minh et al. 2020) on XSEDE (Towns et al. 2014) and CIPRES (Miller et al. 2010) platforms. For model selection, ModelFinder (Kalyaanamoorthy et al. 2017) method was selected. The best model according to Bayesian information criterion (BIC) was WAG+R8 and WAG+R7, for the cyt *c*_554_+ and NrfA+ MSAs, respectively. For generation of branch support values, ultra-fast bootstrap (Hoang et al. 2018) and SH-aLRT (Guindon et al. 2010) statistical methods were used. Confidence values were based on 1000 replications for each methods. For Bayesian inference, phylogeny was reconstructed using MrBayes (version 3.2.7a x86_64) (Ronquist et al. 2012) on XSEDE (Towns et al. 2014) and CIPRES (Miller et al. 2010) platforms. Since MrBayes does not include the FreeRate model for heterogeneity across sites available at the ModelFinder (Kalyaanamoorthy et al. 2017; Minh et al. 2020), the best model was selected using the Smart model selection (SMS) (Lefort et al. 2017) at the ATGC: Montpellier Bioinformatics Platform (http://www.atgc-montpellier.fr/sms/). The best model according to BIC was WAG +G +I for both cyt *c*_554_+ and NrfA+ MSA. Furthermore best-fit parameters for cyt *c*_554_+ MSA were: proportion of invariable sites of 0.035; number of substitution rate categories of 4 and gamma shape parameter of 1.738. For NrfA+ MSA best-fit parameters were proportion of invariable sites of 0.030; number of substitution rate categories of 4; gamma shape parameter of 1.993. Analysis ran using two independent runs with 12 Metropolis-Coupled Markov chain Monte Carlo each for 2 million generations, sampling from the posterior distribution every 5000 generations. Chain convergence was assessed using Tracer version 1.7.2 (Rambaut et al. 2018). Minimum Estimated Sample Size (ESS) and Potential Scale Reduction Factor (PSRF) (Gelman and Rubin 1992) values were within acceptable values. ESS higher than 200 and PSRF between 1.000 and 1.001, respectively. A majority-rule consensus tree with all compatible groups and posterior probabilities of the bipartitions were used to reconstruct the MHC phylogeny, after discarding the first 50% of the sampled trees as burn-in. Tree rooting was performed by the Minimum Ancestral Evolution (MAD) method (Tria et al. 2017). Horizontal gene transfer (HGT) was assessed by comparing the phylogenetic position from the different protein sequences and the classification of the belonging organisms based on NCBI taxonomy database (Federhen 2012). Mismatch in phyla and class positions were considered as an evidence for HGT. Taxonomic classes were assigned to each tip of the maximum likelihood and Bayesian trees previously constructed. Tree visualization and representation was performed using the Interactive Tree Of Life tool (Letunic and Bork 2019).

## Acknowledgments

Financial support was provided by European EC Horizon2020 TIMB3 (Project 810856). Financial support was also provided by Project MOSTMICRO-ITQB with references UIDB/04612/2020 and UIDP/04612/2020. Fundac ão para a Cie ncia e a Tecnologia (FCT) Portugal is also acknowledged for project PTDC/BIA-BQM/4143/2021.

## Notes

### Competing Interest Statement

The authors have declared no competing interest.

