## Supplementary material for "A new paradigm of multiheme cytochrome evolution by grafting and pruning protein modules": suplementary information

### Supplementary tables

**Table S1.** Distance matrix for the N-terminal backbone comparisons. Each PDB code is colored accordingly with the clade where this protein belongs. Distance values are in ångströms and heat mapped in greyscale (lower values are darker).

| | 1FGJ_A | 4N4J_A | 6HIF_A | 4QO5_A | 3SXQ_A | 2OT4_A | 1FT5_A | 4RKM_A | 6I5B_A | 6QVM_A |
|---|---|---|---|---|---|---|---|---|---|---|
| 1FGJ_A | | | | | | | | | | |
| 4N4J_A | 2.239 | | | | | | | | | |
| 6HIF_A | 2.141 | 1.789 | | | | | | | | |
| 4QO5_A | 3.416 | 2.747 | 2.601 | | | | | | | |
| 3SXQ_A | 4.765 | 3.904 | 4.229 | 4.923 | | | | | | |
| 2OT4_A | 4.808 | 3.991 | 4.26 | 4.965 | 0.502 | | | | | |
| 1FT5_A | 5.104 | 5.33 | 6.902 | 4.473 | 5.296 | 5.195 | | | | |
| 4RKM_A | 6.129 | 5.857 | 6.131 | 5.863 | 5.323 | 5.331 | 4.774 | | | |
| 6I5B_A | 5.383 | 4.717 | 4.617 | 4.945 | 4.03 | 4.139 | 4.316 | 4.189 | | |
| 6QVM_A | 5.071 | 4.717 | 4.988 | 5.956 | 4.808 | 4.825 | 3.88 | 4.402 | 2.927 | |
| 1SP3_A | 5.747 | 6.085 | 5.927 | 5.996 | 7.083 | 7.137 | 4.67 | 6.846 | 5.852 | 6.169 |

**Table S2.** Distance matrix for the C-terminal backbone comparisons. Each PDB code is colored accordingly with the clade where this protein belongs. Distance values are in ångströms and heat mapped in greyscale (lower values are darker).

| | 1FGJ_A | 4N4J | 6HIF_A | 4QO5_A | 1FS7_A | 1GU6_A | 1OAH_A | 1QDB_A | 2J7A_A | 3UBR_A | 6V0A_A | 2OT4_A | 3SXQ_A | 4RKM_A | 6I5B_A | 6QVM_A |
|---|---|---|---|---|---|---|---|---|---|---|---|---|---|---|---|---|
| 1FGJ_A | | | | | | | | | | | | | | | | |
| 4N4J | 2.239 | | | | | | | | | | | | | | | |
| 6HIF_A | 2.141 | 1.789 | | | | | | | | | | | | | | |
| 4QO5_A | 3.416 | 2.747 | 2.601 | | | | | | | | | | | | | |
| 1FS7_A | 4.828 | 4.557 | 4.689 | 5.12 | | | | | | | | | | | | |
| 1GU6_A | 4.715 | 4.534 | 4.698 | 4.99 | 1.269 | | | | | | | | | | | |
| 1OAH_A | 4.614 | 4.614 | 5.069 | 4.859 | 2.752 | 2.562 | | | | | | | | | | |
| 1QDB_A | 4.684 | 4.522 | 4.863 | 5.109 | 1.04 | 1.078 | 2.679 | | | | | | | | | |
| 2J7A_A | 4.751 | 4.493 | 5.272 | 5.115 | 2.833 | 2.56 | 1.183 | 2.71 | | | | | | | | |
| 3UBR_A | 4.969 | 4.719 | 4.855 | 5.078 | 1.751 | 0.998 | 2.711 | 1.541 | 2.721 | | | | | | | |
| 6V0A_A | 6.036 | 4.369 | 4.784 | 4.842 | 2.406 | 2.36 | 2.363 | 2.664 | 2.478 | 2.608 | | | | | | |
| 2OT4_A | 4.808 | 3.991 | 4.26 | 4.965 | 2.642 | 2.419 | 2.545 | 2.532 | 2.575 | 2.574 | 2.403 | | | | | |
| 3SXQ_A | 4.765 | 3.904 | 4.229 | 4.923 | 2.615 | 2.388 | 2.564 | 2.48 | 2.505 | 2.547 | 2.373 | 0.502 | | | | |
| 4RKM_A | 6.129 | 5.857 | 6.131 | 5.863 | 4.362 | 6.159 | 4.5 | 4.117 | 4.86 | 4.258 | 4.359 | 5.331 | 5.323 | | | |
| 6I5B_A | 5.383 | 4.717 | 4.617 | 4.945 | 3.292 | 3.023 | 3.137 | 3.229 | 3.694 | 2.941 | 3.482 | 4.139 | 4.03 | 4.189 | | |
| 6QVM_A | 5.071 | 4.717 | 4.988 | 5.956 | 3.931 | 3.894 | 3.955 | 4.491 | 4.159 | 4.015 | 3.523 | 4.825 | 4.808 | 4.402 | 2.927 | |
| 1SP3_A | 5.747 | 6.085 | 5.927 | 5.996 | 6.824 | 6.66 | 6.785 | 6.945 | 6.837 | 6.682 | 6.9 | 7.137 | 7.083 | 6.846 | 5.852 | 6.169 |

**Table S3.** Distance matrix for the N-terminal heme-core comparisons. Each PDB code is colored accordingly with the clade where this protein belongs. Distance values are in ångströms and heat mapped in greyscale (lower values are darker).

| | 1FGJ_A | 4N4J_A | 6HIF_A | 4QO5_A | 2OT4_A | 3SXQ_A | 1FT5_A | 4RKM_A | 6I5B_A | 6QVM_A |
|---|---|---|---|---|---|---|---|---|---|---|
| 1FGJ_A | | | | | | | | | | |
| 4N4J_A | 0.452 | | | | | | | | | |
| 6HIF_A | 0.42 | 0.268 | | | | | | | | |
| 4QO5_A | 0.367 | 0.448 | 0.52 | | | | | | | |
| 2OT4_A | 1.204 | 1.216 | 1.294 | 1.175 | | | | | | |
| 3SXQ_A | 1.227 | 1.267 | 1.328 | 1.218 | 0.193 | | | | | |
| 1FT5_A | 0.818 | 1.064 | 1.068 | 0.872 | 1.039 | 1.003 | | | | |
| 4RKM_A | 0.842 | 1 | 0.98 | 0.943 | 1.012 | 0.974 | 0.626 | | | |
| 6I5B_A | 0.784 | 0.924 | 0.886 | 0.873 | 0.99 | 0.993 | 0.654 | 0.524 | | |
| 6QVM_A | 0.767 | 0.825 | 0.813 | 0.803 | 1.07 | 1.107 | 0.801 | 0.66 | 0.412 | |
| 1SP3_A | 1.372 | 1.623 | 1.617 | 1.451 | 1.326 | 1.263 | 0.967 | 1.137 | 1.154 | 1.472 |

**Table S4.** Distance matrix for the C-terminal heme-core comparisons. Each PDB code is colored accordingly with the clade where this protein belongs. Distance values are in ångströms and heat mapped in greyscale (lower values are darker).

| | 1FGJ_A | 4N4J_A | 6HIF_A | 4QO5_A | 1FS7_A | 1GU6_A | 1OAH_A | 1QDB_A | 2J7A_A | 3UBR_A | 6V0A_A | 2OT4_A | 3SXQ_A | 4RKM_A | 6I5B_A | 6QVM_A |
|---|---|---|---|---|---|---|---|---|---|---|---|---|---|---|---|---|
| 1FGJ_A | | | | | | | | | | | | | | | | |
| 4N4J_A | 0.516 | | | | | | | | | | | | | | | |
| 6HIF_A | 0.34 | 0.417 | | | | | | | | | | | | | | |
| 4QO5_A | 0.653 | 0.569 | 0.649 | | | | | | | | | | | | | |
| 1FS7_A | 1.269 | 1.137 | 1.229 | 1.061 | | | | | | | | | | | | |
| 1GU6_A | 1.094 | 0.983 | 1.047 | 0.909 | 0.321 | | | | | | | | | | | |
| 1OAH_A | 1.358 | 1.187 | 1.318 | 1.121 | 0.344 | 0.489 | | | | | | | | | | |
| 1QDB_A | 1.216 | 1.092 | 1.181 | 1.043 | 0.16 | 0.318 | 0.378 | | | | | | | | | |
| 2J7A_A | 1.335 | 1.195 | 1.298 | 1.147 | 0.382 | 0.503 | 0.235 | 0.385 | | | | | | | | |
| 3UBR_A | 1.174 | 1.083 | 1.134 | 1.048 | 0.333 | 0.324 | 0.474 | 0.314 | 0.438 | | | | | | | |
| 6V0A_A | 1.435 | 1.283 | 1.412 | 1.223 | 0.399 | 0.558 | 0.373 | 0.419 | 0.365 | 0.523 | | | | | | |
| 2OT4_A | 1.114 | 1.016 | 1.083 | 1.046 | 0.515 | 0.488 | 0.592 | 0.45 | 0.52 | 0.467 | 0.557 | | | | | |
| 3SXQ_A | 1.001 | 0.937 | 0.982 | 1.025 | 0.682 | 0.598 | 0.775 | 0.618 | 0.681 | 0.561 | 0.707 | 0.302 | | | | |
| 4RKM_A | 1.614 | 1.447 | 1.646 | 1.372 | 0.917 | 0.956 | 0.872 | 0.941 | 0.909 | 1.025 | 0.815 | 1.118 | 1.202 | | | |
| 6I5B_A | 1.309 | 1.041 | 1.278 | 1.082 | 0.908 | 0.918 | 0.802 | 0.928 | 0.837 | 0.97 | 0.892 | 1.031 | 1.034 | 0.775 | | |
| 6QVM_A | 1.432 | 1.481 | 1.468 | 1.182 | 1.007 | 0.948 | 1.054 | 1.036 | 1.006 | 1.009 | 1.029 | 1.134 | 1.174 | 1.158 | 1.275 | |
| 1SP3_A | 1.813 | 1.925 | 1.846 | 2.107 | 2.312 | 2.152 | 2.435 | 2.283 | 2.433 | 2.231 | 2.46 | 2.277 | 2.186 | 2.266 | 2.253 | 2.432 |

**Table S5.** Dataset collected from NCBI RefSeq database before clustering. Number of sequences, overall mean distance, phyla composition for each group of protein sequences.

| Proteins | Number of sequences | Overall mean distance (± SE) | Phyla |
|---|---|---|---|
| HAO/HDH | 165 | 0.468 ± 0.012 | *Nitrospirae; Planctomycetes; Proteobacteria; Verrucomicrobia* |
| IhOCC | 64 | 0.520 ± 0.013 | *Aquificae; Crenarchaeota; Euryarchaeota; Kryptonia (candidatus); Planctomycetes; Proteobacteria; Thermodesulfobacteria* |
| NrfA | 4887 | 0.503 ±0.012 | *Acidobacteria; Actinobacteria; Bacteroidetes; Balneolaeota; Chloroflexi; Chrysiogenetes; Deinococcus-Thermus; Euryarchaeota; Firmicutes; Ignavibacteriae; Kiritimatiellaeota; Lentisphaerae; Nitrospirae; Planctomycetes; Proteobacteria; Spirochaetes; Verrucomicrobia* |
| ONR | 71 | 0.444 ± 0.012 | *Deferribacteres; Proteobacteria; Thermodesulfobacteria; Nitrospirae* |
| Cyt $c_{554}$ | 95 | 0.351 ± 0.018 | *Proteobacteria; Nitrospirae* |
| MccA | 412 | 0.401 ± 0.011 | *Proteobacteria* |
| OcwA | 2 | 0.010 ± 0.004 | *Firmicutes* |
| OmhA | 4 | 0.173 ± 0.009 | *Firmicutes* |

### Supplementary figure legends

**Fig. S1.** Pairwise alignment for OcwA and OmhA sequences**.** Secondary structure is depicted near each sequence. Black rectangles mark exact residue matches. Similar matches are marked in bold. Heme-binding motifs are underlined in red.

**Fig. S2.** Representation of the hemes and distal axial ligand positions. Diamonds represent the hemes. Yellow diamonds represent catalytic hemes. Hemes are labeled sequentially considering the designations of Table2. Respective distal axial ligands are marked with a prime symbol. Displaced distal axial ligands are in grey with an arrow indicating the change from the common placement to the new position.

**Fig. S3.** Taxonomic classification for clade 1 MHC. Clade 1 subtrees retrieved form the main trees (fig. 3 A – D): (*A*) Maximum likelihood subtree for the cyt $c_{554}$+ MSA; (*B*) Bayesian subtree for the cyt $c_{554}$+ MSA; (*C*) Maximum likelihood subtree for the NrfA+ MSA; (*D*) Bayesian subtree for the NrfA+ MSA. Taxonomic classification to the level of class for the organisms included are presented next to tip of branches or collapsed branches. Nodes with bootstrap/SH-aLTR or posterior probabilities values above 70% contain black dots. Tree scale represents number of substitutions per site.

**Fig. S4.** Taxonomic classification for clade 2 MHC. Clade 2 subtrees retrieved form the main trees (fig. 3 A – D): (*A*) Maximum likelihood subtree for the cyt $c_{554}$+ MSA; (*B*) Bayesian subtree for the cyt $c_{554}$+ MSA; (*C*) Maximum likelihood subtree for the NrfA+ MSA; (*D*) Bayesian subtree for the NrfA+ MSA. Taxonomic classification to the level of class for the organisms included are presented next to tip of branches or collapsed branches. Nodes with bootstrap/SH-aLTR or posterior probabilities values above 70% contain black dots. Tree scale represents number of substitutions per site.

**Fig. S5.** Taxonomic classification for clade 3 MHC. Clade 3 subtrees retrieved form the main trees (fig. 3 a – d): (*A*) Maximum likelihood subtree for the cyt $c_{554}$+ MSA; (*B*) Maximum likelihood subtree for the NrfA+ MSA; (*C*) Bayesian subtree cyt $c_{554}$+ MSA; (*D*) Bayesian subtree for the NrfA+ MSA. Taxonomic classification to the level of class for the organisms included are presented next to tip of branches or collapsed branches. Nodes with bootstrap/SH-aLTR or posterior probabilities values above 70% contain black dots. Tree scale represents number of substitutions per site.

**Fig. S6.** Conservation of the cross-linking tyrosine for Clade 1**.** Subtrees retrieved form the main trees (fig. 3 A – D): (*A*) Maximum likelihood subtree for the cyt $c_{554}$+ MSA; (*B*) Bayesian subtree for the cyt $c_{554}$+ MSA; (*C*) Maximum like likelihood subtree for the NrfA+ MSA; (*D*) Bayesian subtree for the NrfA+ MSA. Nodes with bootstrap/SH-aLTR (ML trees) or posterior probabilities (Bayesian trees) values above 70% contain black dots. Tree scale represents number of substitutions per site.

**Fig. S7.** Conservation of the lysine at the proximal axial ligand position of the catalytic heme for clade 2. Subtrees retrieved form the main trees (fig. 3B and 3D): (*A*) Maximum likelihood subtree for the NrfA+ MSA; (*B*) Bayesian subtree for the NrfA+ MSA. Nodes with bootstrap/SH-aLTR (ML tree) or posterior probabilities (Bayesian tree) values above 70% contain black dots. Tree scale represents number of substitutions per site.

**Fig. S8.** Cladogram of fig. 4 including patterns of heme types. Patterns of heme types are indicated as in Table 2; "o" stands for "Other"; changes of heme types are underlined.

### Supplementary figures

#### Fig. S1

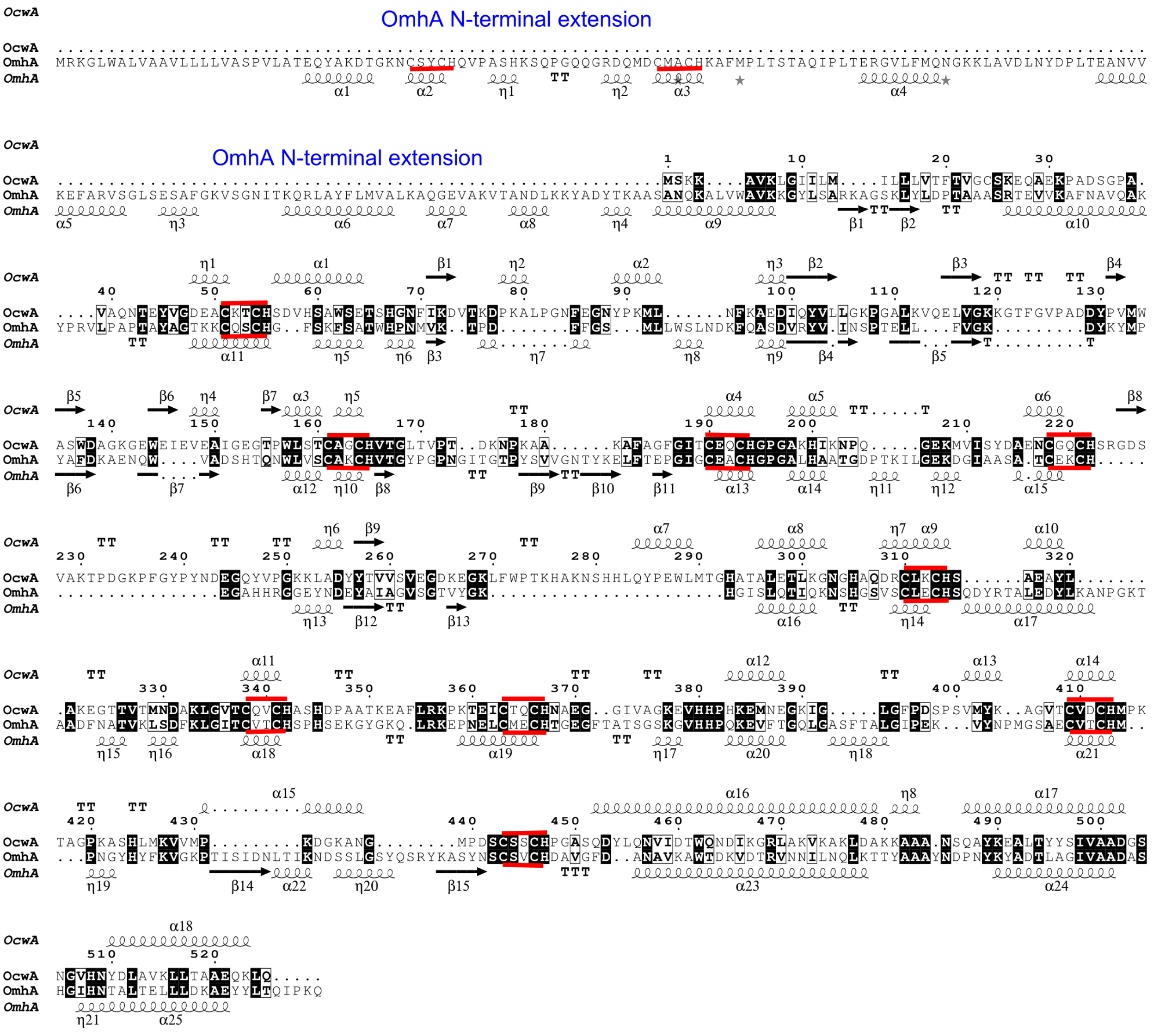

**Fig. S2**

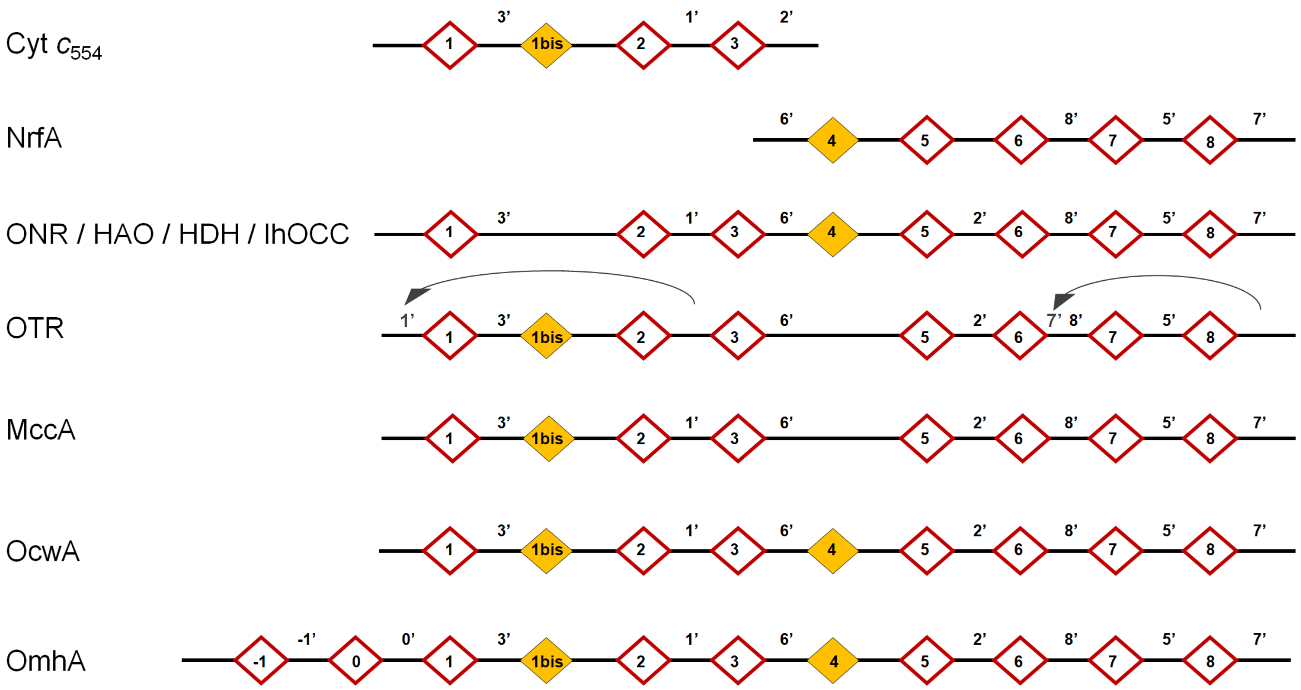

**Fig. S3**

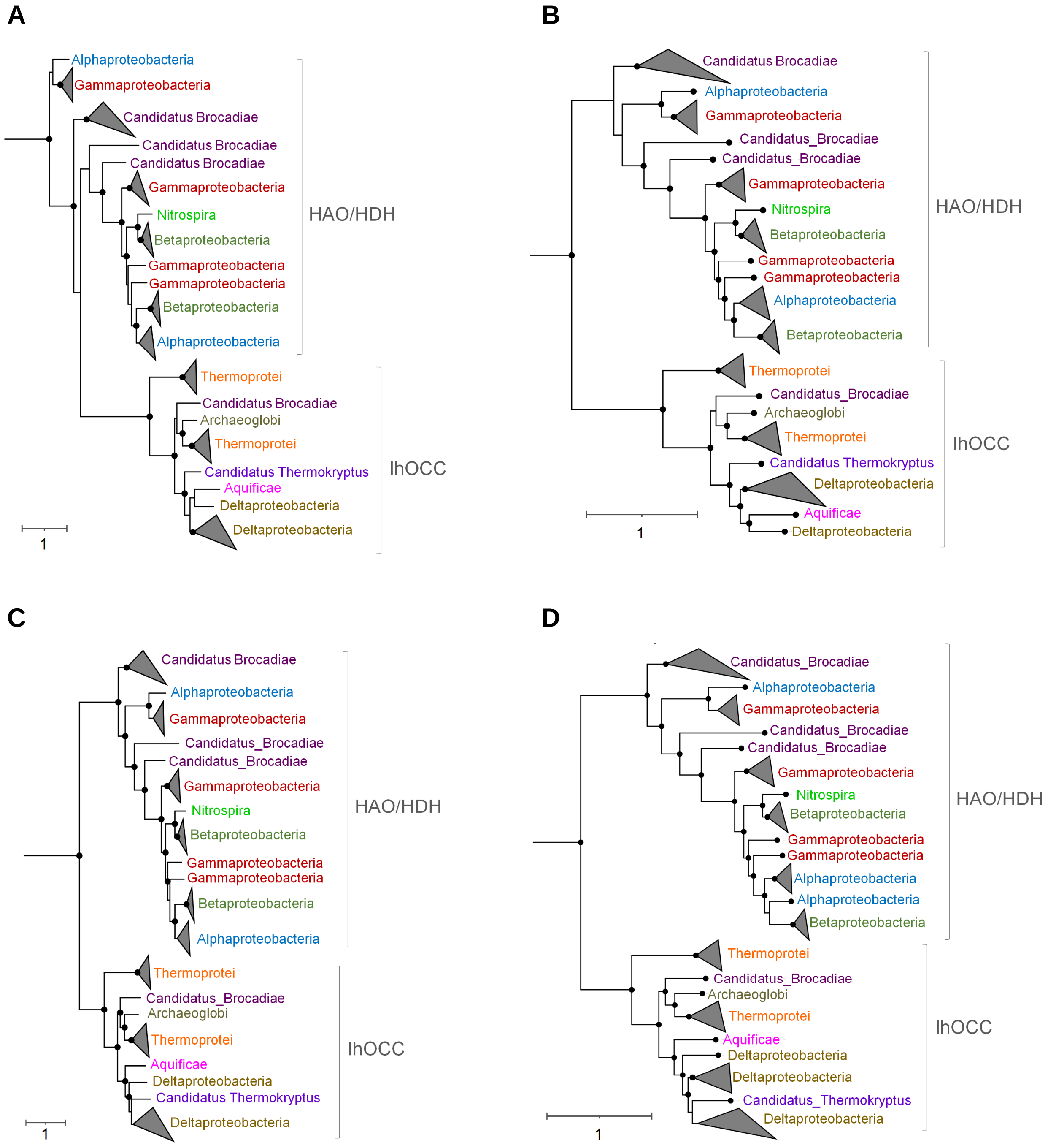
A
Alphaproteobacteria
Gammaproteobacteria
Candidatus Brocadiae
Candidatus Brocadiae
Candidatus Brocadiae
Gammaproteobacteria
Nitrospira
Betaproteobacteria
Gammaproteobacteria
Gammaproteobacteria
Betaproteobacteria
Alphaproteobacteria
HAO/HDH
Thermoprotei
Candidatus Brocadiae
Archaeoglobi
Thermoprotei
Candidatus Thermokryptus
Aquificae
Deltaproteobacteria
Deltaproteobacteria
IhOCC
1
B
Candidatus Brocadiae
Alphaproteobacteria
Gammaproteobacteria
Candidatus_Brocadiae
Candidatus_Brocadiae
Gammaproteobacteria
Nitrospira
Betaproteobacteria
Gammaproteobacteria
Gammaproteobacteria
Alphaproteobacteria
Betaproteobacteria
HAO/HDH
Thermoprotei
Candidatus_Brocadiae
Archaeoglobi
Thermoprotei
Candidatus Thermokryptus
Deltaproteobacteria
Aquificae
Deltaproteobacteria
IhOCC
1
C
Candidatus Brocadiae
Alphaproteobacteria
Gammaproteobacteria
Candidatus_Brocadiae
Candidatus_Brocadiae
Gammaproteobacteria
Nitrospira
Betaproteobacteria
Gammaproteobacteria
Gammaproteobacteria
Betaproteobacteria
Alphaproteobacteria
HAO/HDH
Thermoprotei
Candidatus_Brocadiae
Archaeoglobi
Thermoprotei
Aquificae
Deltaproteobacteria
Candidatus Thermokryptus
Deltaproteobacteria
IhOCC
1
D
Candidatus_Brocadiae
Alphaproteobacteria
Gammaproteobacteria
Candidatus_Brocadiae
Candidatus_Brocadiae
Gammaproteobacteria
Nitrospira
Betaproteobacteria
Gammaproteobacteria
Gammaproteobacteria
Alphaproteobacteria
Alphaproteobacteria
Betaproteobacteria
HAO/HDH
Thermoprotei
Candidatus_Brocadiae
Archaeoglobi
Thermoprotei
Aquificae
Deltaproteobacteria
Deltaproteobacteria
Candidatus_Thermokryptus
Deltaproteobacteria
IhOCC
1

**Fig. S4**

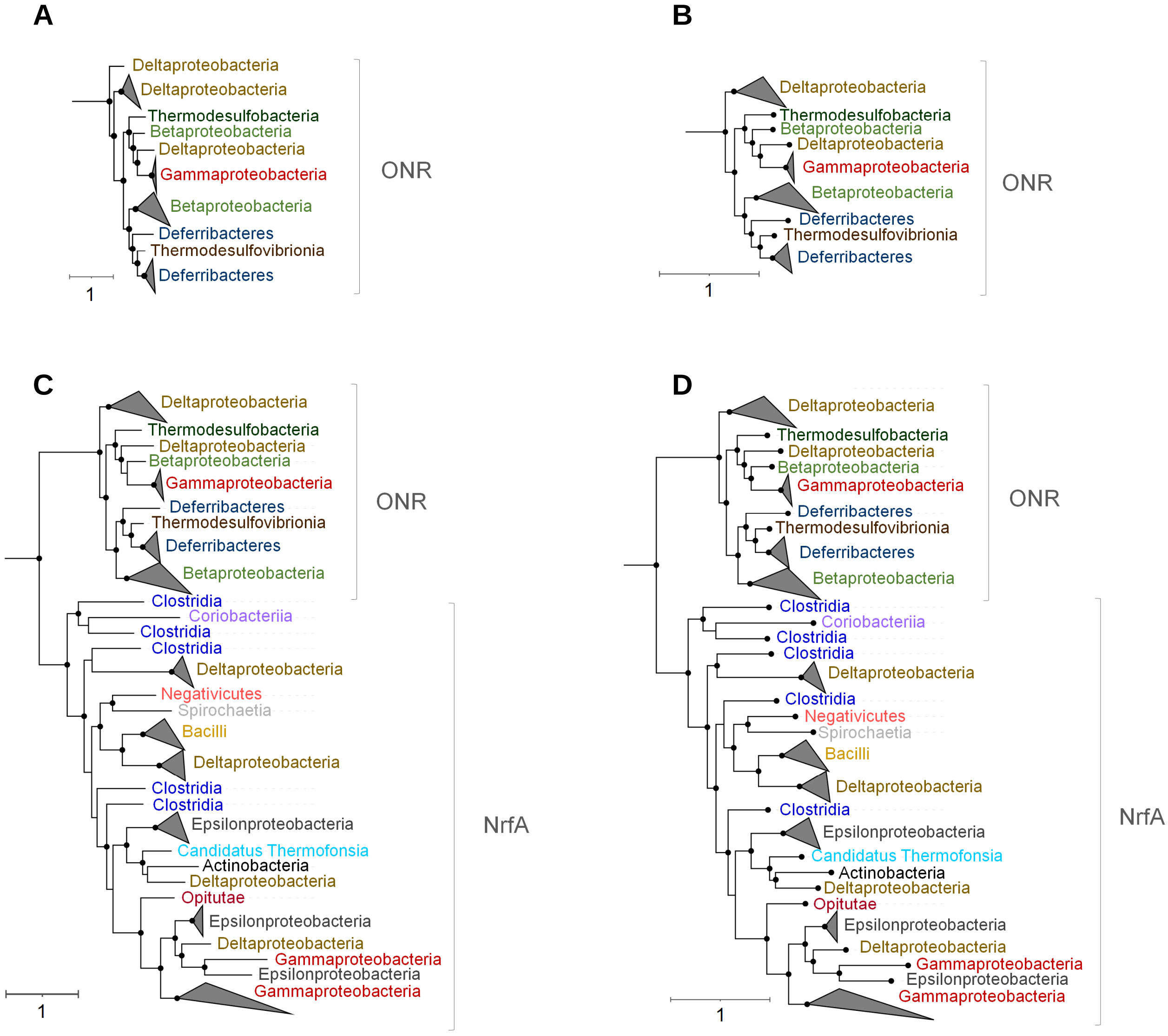

A
Deltaproteobacteria
Deltaproteobacteria
Thermodesulfobacteria
Betaproteobacteria
Deltaproteobacteria
Gammaproteobacteria
Betaproteobacteria
Deferribacteres
Thermodesulfovibrionia
Deferribacteres
ONR
1
B
Deltaproteobacteria
Thermodesulfobacteria
Betaproteobacteria
Deltaproteobacteria
Gammaproteobacteria
Betaproteobacteria
Deferribacteres
Thermodesulfovibrionia
Deferribacteres
ONR
1
C
Deltaproteobacteria
Thermodesulfobacteria
Deltaproteobacteria
Betaproteobacteria
Gammaproteobacteria
Deferribacteres
Thermodesulfovibrionia
Deferribacteres
Betaproteobacteria
ONR
Clostridia
Coriobacteriia
Clostridia
Clostridia
Deltaproteobacteria
Negativicutes
Spirochaetia
Bacilli
Deltaproteobacteria
Clostridia
Clostridia
Epsilonproteobacteria
Candidatus Thermofonsia
Actinobacteria
Deltaproteobacteria
Opitutae
Epsilonproteobacteria
Deltaproteobacteria
Gammaproteobacteria
Epsilonproteobacteria
Gammaproteobacteria
NrfA
1
D
Deltaproteobacteria
Thermodesulfobacteria
Deltaproteobacteria
Betaproteobacteria
Gammaproteobacteria
Deferribacteres
Thermodesulfovibrionia
Deferribacteres
Betaproteobacteria
ONR
Clostridia
Coriobacteriia
Clostridia
Clostridia
Deltaproteobacteria
Clostridia
Negativicutes
Spirochaetia
Bacilli
Deltaproteobacteria
Clostridia
Epsilonproteobacteria
Candidatus Thermofonsia
Actinobacteria
Deltaproteobacteria
Opitutae
Epsilonproteobacteria
Deltaproteobacteria
Gammaproteobacteria
Epsilonproteobacteria
Gammaproteobacteria
NrfA
1

**Fig. S5**

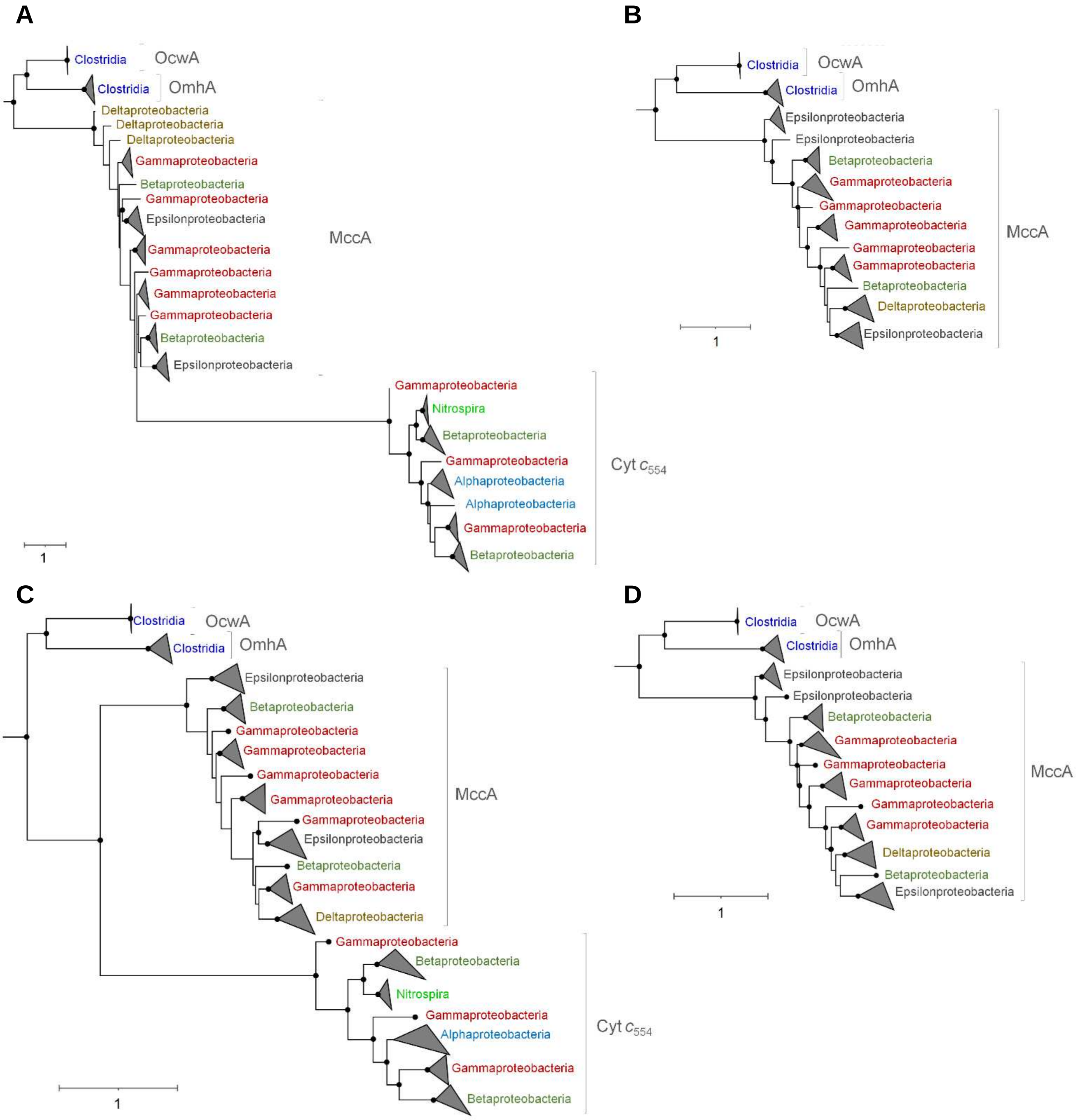
A
Clostridia
OcwA
Clostridia
OmhA
Deltaproteobacteria
Deltaproteobacteria
Deltaproteobacteria
Gammaproteobacteria
Betaproteobacteria
Gammaproteobacteria
Epsilonproteobacteria
MccA
Gammaproteobacteria
Gammaproteobacteria
Gammaproteobacteria
Gammaproteobacteria
Betaproteobacteria
Epsilonproteobacteria
Gammaproteobacteria
Nitrospira
Betaproteobacteria
Gammaproteobacteria
Cyt $c_{554}$
Alphaproteobacteria
Alphaproteobacteria
Gammaproteobacteria
Betaproteobacteria
1
B
Clostridia
OcwA
Clostridia
OmhA
Epsilonproteobacteria
Epsilonproteobacteria
Betaproteobacteria
Gammaproteobacteria
Gammaproteobacteria
Gammaproteobacteria
MccA
Gammaproteobacteria
Gammaproteobacteria
Betaproteobacteria
Deltaproteobacteria
Epsilonproteobacteria
1
C
Clostridia
OcwA
Clostridia
OmhA
Epsilonproteobacteria
Betaproteobacteria
Gammaproteobacteria
Gammaproteobacteria
Gammaproteobacteria
MccA
Gammaproteobacteria
Gammaproteobacteria
Epsilonproteobacteria
Betaproteobacteria
Gammaproteobacteria
Deltaproteobacteria
Gammaproteobacteria
Betaproteobacteria
Nitrospira
Gammaproteobacteria
Alphaproteobacteria
Cyt $c_{554}$
Gammaproteobacteria
Betaproteobacteria
1
D
Clostridia
OcwA
Clostridia
OmhA
Epsilonproteobacteria
Epsilonproteobacteria
Betaproteobacteria
Gammaproteobacteria
Gammaproteobacteria
MccA
Gammaproteobacteria
Gammaproteobacteria
Gammaproteobacteria
Deltaproteobacteria
Betaproteobacteria
Epsilonproteobacteria
1

Fig. S6

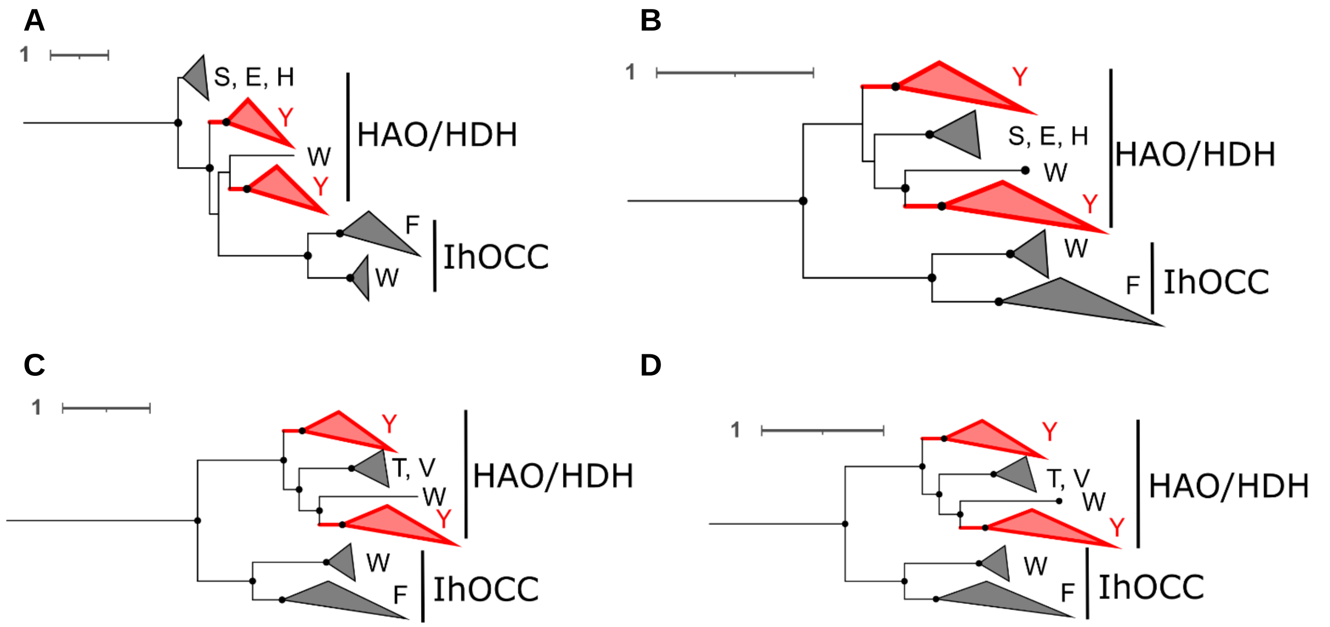
A
1
S, E, H
Y
W
Y
HAO/HDH
F
W
IhOCC
B
1
Y
S, E, H
W
Y
HAO/HDH
W
F
IhOCC
C
1
Y
T, V
W
Y
HAO/HDH
W
F
IhOCC
D
1
Y
T, V
W
Y
HAO/HDH
W
F
IhOCC

Fig. S7

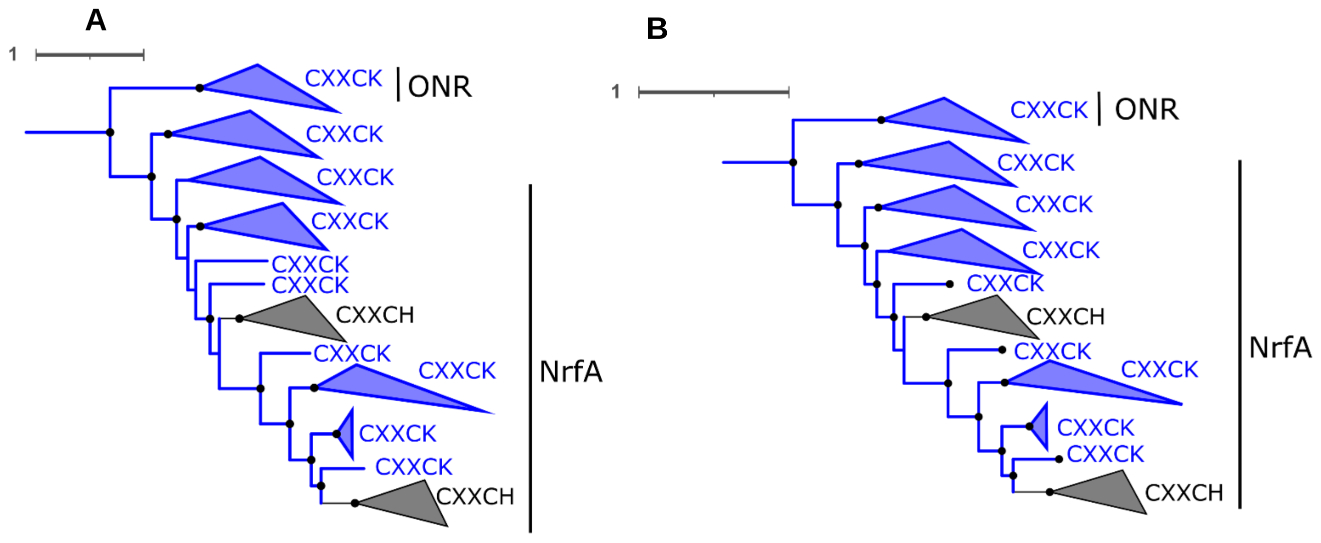
A
1
CXXCK
ONR
CXXCK
CXXCK
CXXCK
CXXCK
CXXCK
CXXCH
CXXCK
CXXCK
CXXCK
CXXCK
CXXCH
NrfA
B
1
CXXCK
ONR
CXXCK
CXXCK
CXXCK
CXXCK
CXXCH
CXXCK
CXXCK
CXXCK
CXXCK
CXXCH
NrfA

**Fig. S8**

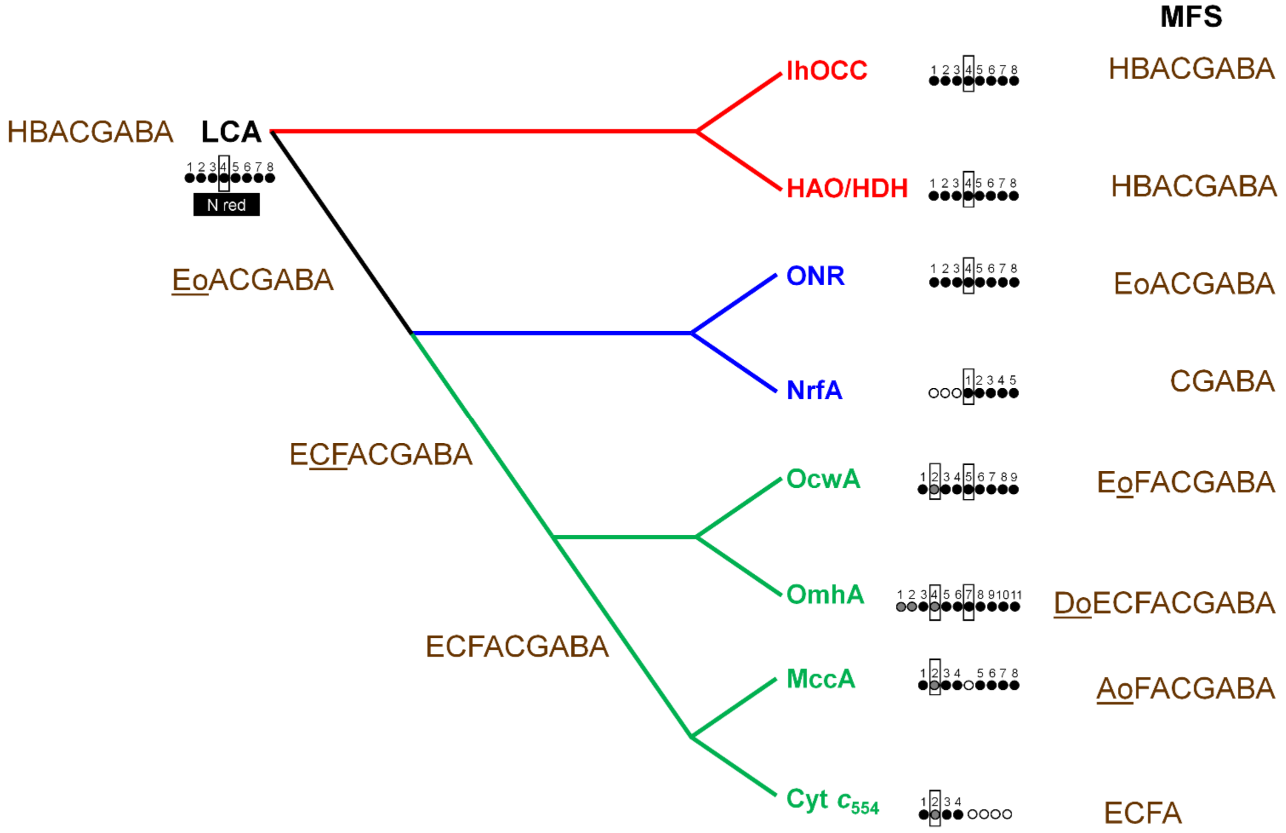
MFS
IhOCC
HBACGABA
HBACGABA
LCA
N red
HAO/HDH
HBACGABA
EoACGABA
ONR
EoACGABA
NrfA
CGABA
ECFACGABA
OcwA
EoFACGABA
OmhA
DoECFACGABA
ECFACGABA
MccA
AoFACGABA
Cyt c554
ECFA
